# On a non-standard two-species stochastic competing system and a related degenerate parabolic equation

**DOI:** 10.1101/2020.03.21.001347

**Authors:** H. Yoshioka, Y. Yoshioka

## Abstract

We propose and analyse a new stochastic competing two-species population dynamics model. Competing algae population dynamics in river environment, important engineering problems, motivate this model. It is a system of stochastic differential equations (SDEs) and has a characteristic that the two populations are competing with each other through the environmental capacities. Unique existence of the uniformly bounded strong solution is proven and an attractor is identified. The Kolmogorov backward equation associated with the population dynamics is formulated and its unique solvability in a Banach space with a weighted norm is discussed. Our mathematical analysis results can be effectively utilized for a base-stone of modelling, analysis, and control of the competing population dynamics.

## 1 Introduction

Competing population dynamics have long been a central topic in mathematical biology. Especially, stochastic process models based on stochastic differential equations (SDEs) [13] have served as efficient mathematical tools for modelling and analysis of population dynamics.

There exist a vast number of researches on competing population dynamics models. Predator–prey systems under stochastic environment have rich mathematical structures, and have been studied in detail [5]. Models with degenerate drift and/or diffusion coefficients are often realistic candidates of stochastic dynamical systems especially in ecology and epidemiology. Lv et al. [11] showed that solutions to a stochastic Lotka–Volterra model having degenerate diffusion coefficients are almost surely confined in a compact set. A stochastic epidemic model having a saturated contact rate has been studied with an SDE possessing degenerate drift and diffusion coefficients [9]. Grandits et al. [6] analysed the value of information in epidemiological dynamics using a system of SDEs having degenerate coefficients combined with a dynamic programming principle. Cai et al. [2, 1] considered a system of SDEs driven by independent and correlated Brownian noises multiplied by degenerate diffusion coefficients.

In this paper, we propose a new stochastic two-species competing population dynamics model as a system of SDEs. The model is motivated by competing population dynamics of filamentous and non-filamentous algae attached on the riverbed in dam-downstream reaches [17]. The nuisance filamentous algae, such as the periphyton *Cladophora glomerata*, are weak against turbulent river flows found in natural rivers, but can persist under low-flow conditions occurring in rivers where humans regulate the flow conditions through dam operations. In addition, aquatic species like *Plecoglossus altivelis altivelis* eat but cannot digest them [18]. On the other hand, non-filamentous algae, such as diatoms serving as staple foods of the fish, are more tolerant against turbulence but possibly have smaller intrinsic growth rates (Schmidt et al. [16] for periphyton and Morin et al. [12] for diatoms.). They are competing on the riverbed, and tracking and predicting their population dynamics are important industrial problem; however, least attention has been made on modelling and analysis of the population dynamics except for the engineering model [17].

Our first contribution in this paper is mathematical analysis on the population dynamics model. Our system of SDEs, which is new to the best of our knowledge, has a characteristic that the species are interacting with each other through the environmental capacities. We show that this formulation is well-posed and the system admits a unique bounded strong solution. This is a fundamentally important contribution because guaranteeing well-posedness of a mathematical model is essential in its analysis and computation. We also show that the solution eventually converges toward a part of the boundary of a compact set under certain conditions. Our another contribution is a unique solvability result of the Kolmogorov backward equation (KBE) associated with the population dynamics. The KBE is a fundamental equation when statistically assessing the population dynamics. This equation is of a degenerate parabolic type and does not always have classical solutions satisfying the equation in the point-wise sense [14]. With the help of the function analysis techniques for degenerate parabolic problems (e.g., see [8, 7]), we show that the KBE has a variational weak solution in a Banach space with a weighted norm. Throughout this paper, we set 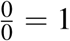 for the sake of brevity of analysis.

## 2 Population dynamics

### 2.1 System dynamics

We consider continuous-time two-species population dynamics in a habitat. The time is denoted as *t* ≥ 0. There are two species in the model and their populations are denoted as *X*_*t*_ and *Y*_*t*_ at time *t*, respectively. We assume that the populations are living in a habitat with the unit area, and that the total populations represent the areas shared by the populations. Namely, we assume 0 ≤ *X*_*t*_, *Y*_*t*_, *X*_*t*_ + *Y*_*t*_ ≤ 1. Accordingly, we set the compact triangular domain *A* and its interior *Â* as *A* = {(*x, y*); 0 ≤ *x, y, x* + *y*≤ 1} and *Â* = {(*x, y*); 0 < *x, y, x* + *y* < 1}, respectively. The boundary of *A* is denoted as *∂A* = *A− Â*. We do not consider spatially-distributed population dynamics, but will be considered elsewhere. Hereafter, the following elementary inequality is used:

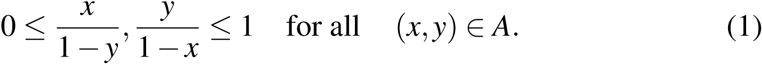

The system of SDEs governing the two-species population dynamics is proposed as follows:

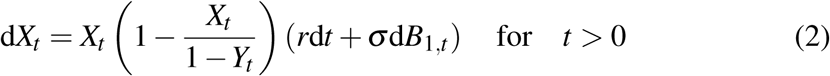

and

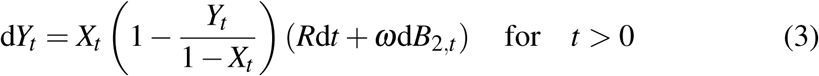

subject to an initial condition (*X*_0_,*Y*_0_), where *r, R, σ, ω* > 0 and E[d*B*_1,*t*_ d*B*_2,*t*_] = *ρ*d*t*. We assume |*ρ*| ≤ 1. In (2)-(3), the terms involving d*t* represent deterministic growth, while those involving d*B*_*i,t*_ (*i* = 1, 2) represent the stochastic fluctuations of the dynamics that are potentially correlated. A characteristic of this system is that unit area is shared by the two populations; increasing one population decreases the other’s environmental capacity. The model can be formally considered as a competing population dynamics model having the stochastic growth rates 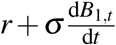 and 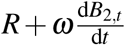 with the state-dependent environmental capacities. Considering the problem background, it is biologically reasonable to assume *r* > *R* and *σ* < *ω*. However, we do not employ this assumption and carry out mathematical analysis under more general conditions.

### 2.2 Mathematical analysis

A fundamental issue of the system (2)-(3) is its unique solvability. The following proposition resolves this issue, guaranteeing that the system is well-posed and the population is certainly confined in *A* as desired. Hereafter, set

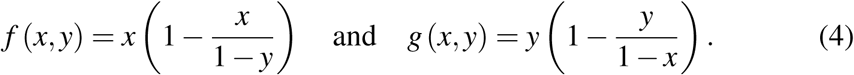

They are Lipschitz continuous in *A*.

#### Proposition 1.

*We have* (*X*_*t*_, *Y*_*t*_) ∈ *A for t* > 0 *a.s., if* (*X*_0_,*Y*_0_) ∈ *A.*

**Proof:** The key of the proof is the Ito’s formula [13]: for sufficiently smooth *h* = *h* (*X*_*t*_, *Y*_*t*_), Set *h*_1_ = ln *X*_*t*_, *h*_2_ = ln*Y*_*t*_, and *h*_3_ = ln(1 *−X*_*t*_ *−Y*_*t*_). We have 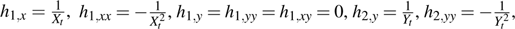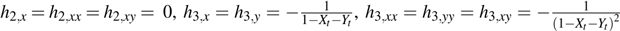. Set the stopping time

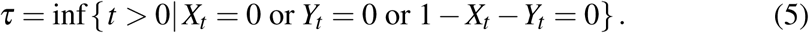

We firstly show *τ* → +∞ a.s. if (*X*_0_,*Y*_0_) *Â*, with which we get (*X*_*t*_, *Y*_*t*_) ∈ *A* a.s. *t* > 0.

There exist the six possible cases: 1. *X*_*τ*_ > 0, *Y*_*τ*_ = 0, 1 − *X*_*τ*_ − *Y*_*τ*_ = 0, 2. *X*_*τ*_ = 0, *Y*_*τ*_ > 0, 1 − *X*_*τ*_ − *Y*_*τ*_ = 0, 3. *X*_*τ*_ = 0, *Y*_*τ*_ = 0, 1 − *X*_*τ*_ − *Y*_*τ*_ > 0, 4. *X*_*τ*_ > 0, *Y*_*τ*_ > 0, 1 − *X*_*τ*_ − *Y*_*τ*_ = 0, 5. *X*_*τ*_ = 0, *Y*_*τ*_ > 0, 1− *X*_*τ*_ − *Y*_*τ*_ > 0, 6. *X*_*τ*_ > 0, *Y*_*τ*_ = 0, 1 − *X*_*τ*_ − *Y*_*τ*_ > 0. Assume (*X*_0_,*Y*_0_) ∈ *Â*. By the symmetry, it is enough to show that the cases 1, 3, 4, 5 do not occur. Now, we have

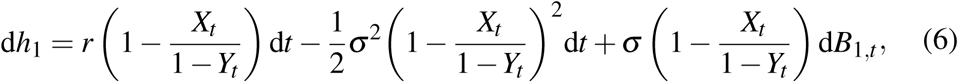

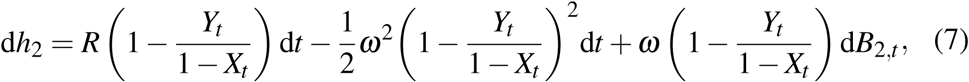

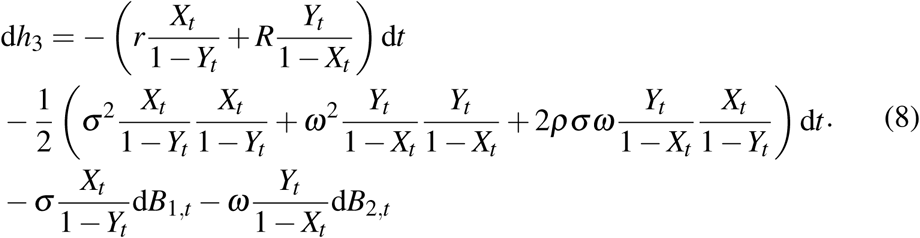

All the coefficients multiplies by d*t*, d*B*_1,*t*_, d*B*_2,*t*_ are bounded for 0 < *t* < *τ* by (1) and (5), while at least one of the light-hand sides of (6)-(8) diverges as *t → τ* − 0 by (5). This is a contradiction if *τ* < +∞, meaning that *τ* = +∞. Notice that the drift and diffusion coefficients can be extended to a globally Lipschitz continuous function over ℝ^2^.

If (*X*_0_,*Y*_0_) ∈ *∂A*, then we get a unique strong solution with the extended coefficients. This completes the proof since (*X*_*t*_, *Y*_*t*_) = (*X*_0_,*Y*_0_) a.s. for *t* > 0. ♠

The next proposition is on a long-time limit of the population dynamics. Set *L* = {(*x, y*); *x, y* ≥ 0, *x* + *y* = 1}, which is a part of *∂A*.

#### Proposition 2.

*If* (*X*_0_,*Y*_0_) ∈ *Â and* 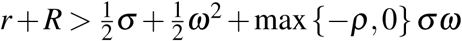, *then a.s.* 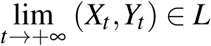.

**Proof:** Recall the conventional iterated logarithm estimate for a 1–D standard Brownian motion *B*_*t*_ (Theorem 5.1.2 of [9]): we have, a.s.

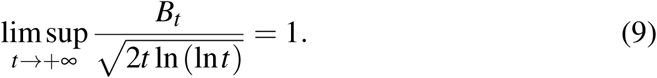

Assume (*X*_0_,*Y*_0_) ∈ *Â*. Then, by the Ito’s formula [13], we get

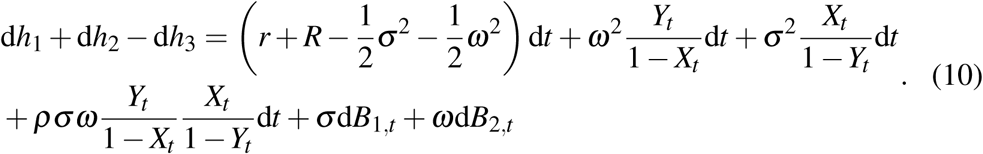

By the assumption of the proposition, if *ρ* < 0, then

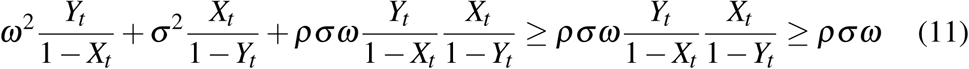

and thus

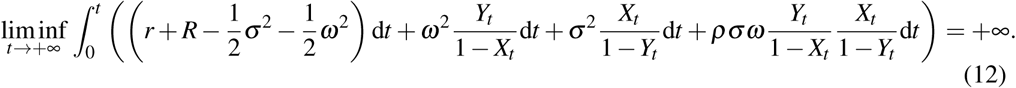

Temporally integrating (6) in (0, *t*) with *t* > 0 and taking 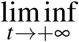 of its both sides get 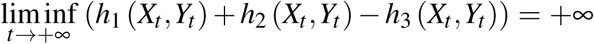. By Proposition 1, this means 1 *−X*_*t*_ +*Y*_*t*_ → 0 as *t* → +∞. The case *ρ* ≥ 0 is essentially the same, and is omitted. ♠

An implication of this proposition is that the both populations do not become extinct if the sum of their growth rates is sufficiently large. It is interesting to see that the correlation *ρ* directly affect the population dynamics.

## 3 Kolmogorov backward equation

### 3.1 Formulation

For each (*t, x, y*) ∈ Ω = [0, *T*] × *A* with *T* > 0, the conditional expectation

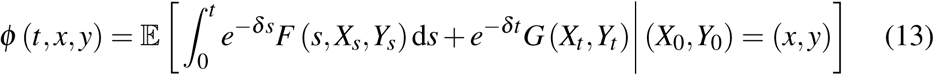

with a discount rate *δ* > 0 and sufficiently regular *F, G* (we assume that they are essentially bounded and continuously differentiable for the sake of simplicity of analysis) statistically evaluates the population dynamics [13], and is therefore an important statistical index. The formal governing equation of *ϕ* is the KBE

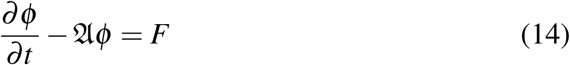

subject to the initial condition *ϕ*_*t*=0_ = *G* in *A*, where

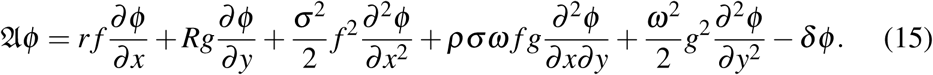

Due to the degenerate coefficients, no boundary condition is necessary along *∂A*. The KBE is degenerate parabolic, meaning that its unique solvability is not a trivial issue, and that there is no guarantee to have solutions satisfying the equation classical point-wise manner. This issue is briefly analysed from a variational viewpoint. Hereafter, we assume *F* = *G* = 0 on *∂A* without significant loss of generality because of the linearity of the KBE.

### 3.2 Mathematical analysis

We introduce a weighted Sobolev space *V* specialized for our problem:

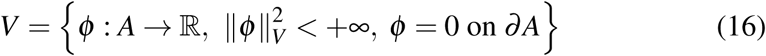

with the norm

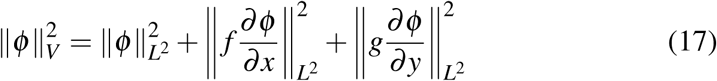

The dual *V*\* of *V* is equipped with the dual norm

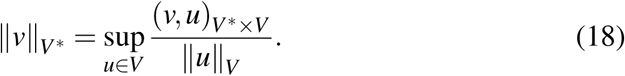

Set the bilinear form *a* : *V* × *V* → ℝ corresponding to the KBE as

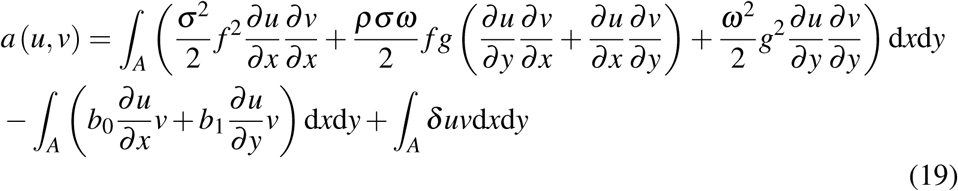

for *u, v* ∈ *V*, where *b* = (*b*_0_, *b*_1_) is given by

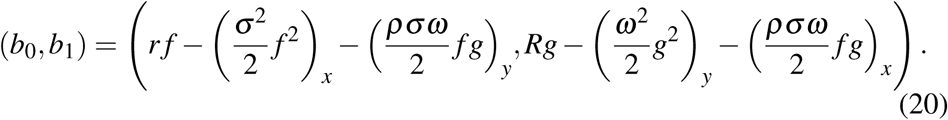

Notice that *b* ⊥ *n* a.e. on *∂A*, where *n* is the outer unit normal on *∂A*.

The next proposition is our main result on the KBE.

#### Proposition 3.

*If* |*ρ*| < 1, *the KBE is uniquely solvable in L*^2^ ((0, *T*); *V*) ∩ *H*^1^ ((0, *T*); *V*\*).

**Proof:** We only present the sketch of the proof because it is sufficient to check the continuity of *a* and the Garding inequality

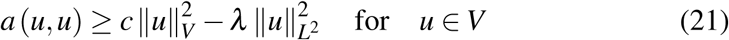

with some constants *c, λ* > 0. For example, see Chapter 3, Section 4.4, Theorem 4.1 of Lions and Magenes [10]. It is sufficient to check these conditions for every functions in 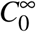 (*A*) because it is dense in *V*. A lengthy but straightforward calculation with integration by parts gives that the Garding inequality is satisfied with the constants 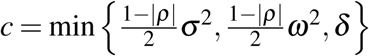 and 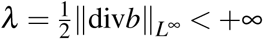. Checking the continuity of *a* is rather straightforward with the Cauchy-Schwartz inequality and is omitted here. ♠

As implied in the proof, the case | *ρ* | = 1 is not covered by the Garding inequality. This case may be handled in the framework of viscosity solutions [4]. It should be noted that using the weighted Sobolev space is essential because of the degenerate diffusion coefficients.

## 4 Conclusions

We proposed a new stochastic population dynamics model and analysed its unique solvability and boundedness. Unique solvability of the associated KBE was also proven in a variational sense. The analysis results suggest that the presented model would potentially serve as a foundation for modelling, analysing, and controlling the algae population dynamics.

There exist several issues to achieve an establishment of a mathematical tool for such purposes. The assumptions made in each proposition should be analysed to see it can be weaken or not. The model parameters have to be identified in natural environment; a part of them is available in the literature. Water quality dynamics, like dissolved silica and dissolved oxygen dynamics, can significantly affect the population dynamics [3]. A key practical issue is that it is often difficult to accurately identify the model parameters through field surveys. Therefore, the model uncertainty should be considered when operating the model. The uncertainty issue can be handled in the framework of the non-linear expectation [15], which enable us to analyse SDEs having uncertain coefficients in a rigorous manner. For example, we can allow for uncertainties in the growth rates and/or correlation coefficient. The presented model does not consider catastrophic flood events that suddenly remove the algae population from the riverbed. Such events can be considered to follow some jump processes. Studying a discrete-time counterpart of the population dynamics is an interesting topic as well. Finally, the available habitat of the algae populations would have seasonal variations according to the river flows, motivating us to consider time-dependent environmental capacities.

## Acknowledgements

JSPS Research Grant No. 19H03073, Kurita Water and Environment Foundation Grant No. 19B018, a grant for ecological survey of a life history of the landlocked ayu *Plecoglossus altivelis altivelis* from the MLIT of Japan, and a research grant for young researchers in Shimane University support this research. The first author engaged this research as a member of Fisheries Ecosystem Project Centre of Shimane University. The authors thank Dr. Zlatko Jovanoski and his colleagues for their valuable comments and suggestions.

## References

[1] S. Cai, Y. Cai, and X. Mao. “A stochastic differential equation SIS epidemic model with two correlated Brownian motions”. In: Nonlinear Dynamics 97.4 (2019), pp. 2175–2187. DOI: 10.1007/s11071-019-05114-2 (cit. on p. 2).

[2] S. Cai, Y. Cai, and X. Mao. “A stochastic differential equation SIS epidemic model with two independent Brownian motions”. In: Journal of Mathematical Analysis and Applications 474.2 (2019), pp. 1536–1550. DOI: 10.1016/j.jmaa.2019.02.039 (cit. on p. 2).

[3] U. Callies, M. Scharfe, and M. Ratto. “Calibration and uncertainty analysis of a simple model of silica-limited diatom growth in the Elbe River”. In: Ecological Modelling 213.2 (2008), pp. 229–244. DOI: 10.1016/j.ecolmodel.2007.12.015 (cit. on p. 8).

[4] M. G. Crandall, H. Ishii, and P. L. Lions. “User’s guide to viscosity solutions of second order partial differential equations”. In: Bulletin of the American Mathematical Society 27.1 (1992), pp. 229–244. DOI: 10.1090/S0273-0979-1992-00266-5 (cit. on p. 7).

[5] N. H. Du and V. H. Sam. “Dynamics of a stochastic LotkaVolterra model perturbed by white noise”. In: Journal of Mathematical Analysis and Applications 324.1 (2006), pp. 82–97. DOI: 10.1016/j.jmaa.2005.11.064 (cit. on p. 2).

[6] P. Grandits, R. M. Kovacevic, and V. M. Veliov. “Optimal control and the value of information for a stochastic epidemiological SIS model”. In: Journal of Mathematical Analysis and Applications 476.2 (2019), pp. 665–695. DOI: 10.1016/j.jmaa.2019.04.005 (cit. on p. 2).

[7] B. Horvath and O. Reichmann. “Dirichlet forms and finite element methods for the SABR model”. In: SIAM Journal on Financial Mathematics 9.2 (2018), pp. 716–754. DOI: 10.1137/16M1066117 (cit. on p. 3).

[8] J. Hozman and T. Tichy. “DG framework for pricing European options under one-factor stochastic volatility models”. In: Journal of Computational and Applied Mathematics 344 (2018), pp. 585–600. DOI: 10.1016/j.cam.2018.05.064 (cit. on p. 3).

[9] G. Lan et al. “A stochastic SIS epidemic model with saturating contact rate”. In: Physica A: Statistical Mechanics and its Applications 529.121504 (2019), pp. 1–14. DOI: 10.1016/j.physa.2019.121504 (cit. on pp. 2, 5).

[10] J. L. Lions and E. Magenes. Non-homogeneous Boundary Value Problems and Applications (Vol. 1). Springer Berlin Heidelberg, 1972 (cit. on p. 7).

[11] J. Lv, X. Zou, and L. Tian. “A geometric method for asymptotic properties of the stochastic Lotka–Volterra model”. In: Communications in Nonlinear Science and Numerical Simulation 67 (2019), pp. 449–459. DOI: 10.1016/j.cnsns.2018.06.031 (cit. on p. 2).

[12] S. Morin, M. Coste, and F. Delmas. “A comparison of specific growth rates of periphytic diatoms of varying cell size under laboratory and field conditions”. In: Hydrobiologia 614.1 (2008), pp. 285–297. DOI: 10.1007/s10750-008-9513-y (cit. on p. 2).

[13] B. Øksendal. Stochastic Differential Equations. Springer Berlin Heidelberg, 2003 (cit. on pp. 2, 4–6).

[14] O. Oleinik and E. V. Radkevic. Second-order Equations with Nonnegative Characteristic Form. Springer Boston, 1973 (cit. on p. 3).

[15] S. Peng. Nonlinear Expectations and Stochastic Calculus under Uncertainty: with Robust CLT and G–Brownian Motion. Springer-Verlag Berlin Heidelberg, 2019 (cit. on p. 8).

[16] T. S. Schmidt et al. “Benthic algal (periphyton) growth rates in response to nitrogen and phosphorus: parameter estimation for water quality models”. In: JAWRA Journal of the American Water Resources Association (2019). DOI: 10.1111/1752-1688.12797 (cit. on p. 2).

[17] Y. Toda and T. Tsujimoto. “Numerical modeling of interspecific competition between filamentous and nonfilamentous periphyton on a flat channel bed”. In: Landscape and Ecological Engineering 6.1 (2010), pp. 81–88. DOI: 10.1007/s11355-009-0093-4 (cit. on p. 2).

[18] H. Yoshioka et al. “Optimal harvesting policy of an inland fishery resource under incomplete information”. In: Applied Stochastic Models in Business and Industry 35.4 (2019), pp. 939–962. DOI: 10.1002/asmb.2428 (cit. on p. 2).

